# Examining the Atypical Structure Adopted by the 37-51 Segment of α-synuclein in the Early Stage of Aggregation

**DOI:** 10.64898/2026.10.19.760142

**Authors:** Adrien Guzzo, Patrice Delarue, Khatuna Kachlishvili, Adrien Nicolaï, Patrick Senet, Gia G. Maisuradze

## Abstract

The aggregation of α-synuclein into insoluble filaments is a hallmark of Parkinson’s disease. Upon membrane binding, α-synuclein adopts a helical conformation, whereas during neurodegeneration it misfolds and aggregates into β-sheet-rich structures. However, the factors that trigger the misfolding and aggregation remain poorly understood. In this study, coarse-grained molecular dynamics simulations are used to characterize the structural ensembles of α-synuclein monomers and dimers at atomic resolution during the early stage of aggregation, covering an effective timescale of 29.7 ms. The simulations reveal that a short sequence spanning residues 37-51 predominantly adopts three distinct conformations: β-hairpin, braid, and random coil. Here, focus is placed on a braid conformation, which, to the best of our knowledge, has not been reported previously in either experimental or computational studies. Although we could not definitively rule out the possibility that the coarse-grained force field overestimates the stability of the braid, two all-atom simulations demonstrated that this structure is sterically feasible and metastable. Furthermore, in monomers, the braid appears to hinder the formation of long-range intramolecular interactions, leaving the N-terminus, NAC (non-amyloid-β component) region, and C-terminus solvent-exposed and potentially facilitating fibril nucleation. In dimers, however, the braid appears to impede intermolecular contacts, especially homotypic intermolecular interactions, thereby slowing aggregation. Upon unfolding of the braid structure, intermolecular contacts, including homotypic interactions, increase noticeably.

## Introduction

α-Synuclein (αS), a small (140 residue) intrinsically disordered protein (IDP), is the primary protein component of Lewy bodies^1^, the insoluble aggregates that are a hallmark of Parkinson’s disease, the second most common neurodegenerative disorder. The precise functions of αS remain elusive. The vast majority of experiments provide strong evidence that αS exists predominantly as an intrinsically disordered monomer in solution^2,3^. However, some experiments suggest that αS can adopt a folded tetrameric state with relatively high helical content under physiological conditions, which is indispensable for normal αS homeostasis^4,5^. Moreover, it adopts a helical conformation upon membrane binding and misfolds and aggregates as a β-sheet in neural degeneration. Nevertheless, the factors that trigger the original misfolding remain unknown.

*In vitro* studies have demonstrated that αS fibrillation proceeds through oligomeric intermediates of diverse sizes and morphologies^6^ and that these oligomeric species are, in fact, more pathogenic than the mature fibrils themselves^7^, high-resolution structures of which have only recently became available^8–12^. Therefore, conformational analysis of these oligomeric intermediates is crucial for understanding the molecular mechanisms underlying fibril formation. Because of the inability to identify and follow the low concentration of oligomers over time experimentally, it is difficult to determine the fundamental microscopic reactions taking place during aggregation^13^. All-atom molecular dynamics (MD) simulation is a valuable tool for studying the details of fibril formation; however, the timescales over which fibrils form limit its usefulness. This problem can be surmounted by using coarse-grained (CG) models coupled with sampling enhancing techniques such as adaptive MD sampling, replica-exchange molecular dynamics (REMD)^14^ or umbrella sampling^15^. For this purpose, use is made of a physics-based UNited-RESidue (UNRES) force field^16,17,18^, which enables to run MD simulations of large proteins in the millisecond timescale and bridge theoretical and experimental worlds. It has been demonstrated that the UNRES can accurately predict not only protein folding kinetics^19,20,21,22,23,24^ but also the mechanisms and crucial sites involved in amyloid fibril formation^25,26,27,28^.

Recently, using coarse-grained MD trajectories, generated with the UNRES force field, we sampled the conformational space of wild type (WT) and mutant (A30P, A53T, and E46K) α-synuclein monomers^29^, and studied the formation of dimers of these systems^30^ over an effective timescale of 29.7 ms. This timescale corresponds to the early stage of aggregation, which still remains poorly understood and is hypothesized to be the most toxic phase of the aggregation process. In these studies, we have detected that a short fragment encompassing region of 37-51 residues has a propensity to form structures that, to the best of our knowledge, have never been observed before experimentally. It should be noted, however, that one of these structures has previously been reported in computational study. More precisely, Yu *et al.*^31^ carried out MD simulations for the WT, and A30P, A53T mutants of α-synuclein and observed that a short segment encompassing region 38-53 reveals a high propensity of forming a b-hairpin (Fig. 1 in Ref. 31); plus, a b-hairpin forms much faster in the mutants than in the WT. Moreover, it has been shown that a b-hairpin interacts with C-terminus, and consequently reduces b-hairpin accessibility for inter-molecular interactions. This finding indicates that if a formation of b-hairpin is a crucial part for αS aggregation, then C-terminus decelerates aggregation process of the system, which is in agreement with experiment^32,33,34^. Therefore, the authors concluded that a b-hairpin formed in region 38-53 is a key player in αS aggregation process^31^. In contrast with these results, by performing REMD with an all-atom force field and explicit solvent for the 63 amino acid residues (35−97) segment of αS, Jain *et al.*^35^ illustrated that the conformations adopted within the 38-53 region are predominantly α-helical.

**Figure 1.**
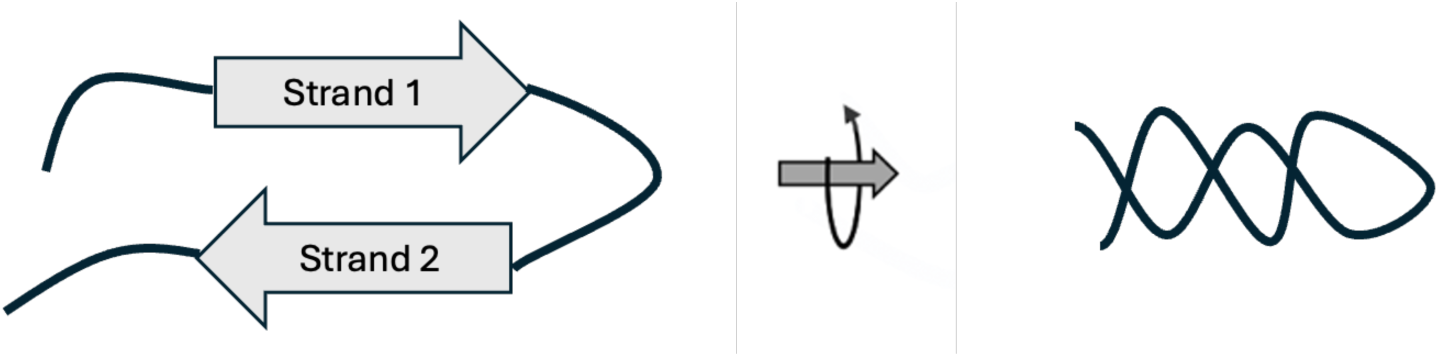
Schematic representation of the braid structure.

Herein, we present a study of a second structural motif detected within residues 37-51 that, to the best of our knowledge, has never been reported before neither experimentally nor computationally. This structure resembles the two strands of a hairpin intertwined through rotation around the hairpin axis; therefore, we termed this motif a *“braid”* (see the schematic representation in Figure 1). The objectives of this paper are to: (i) characterize the structural features of the braid motif; (ii) elucidate a mechanism of formation of the braid motif; (iii) demonstrate that the formation of a braid structure is not an artifact of the coarse-grained force field; and (iv) investigate the influence of the braid structure on the early stages of nucleation process.

## Materials and Methods

### MD Simulations Using UNRES Model

Simulations were carried out using the UNRES force field^16,17,18^. Details of the force field and its parametrization are available in Refs. 14,16-24 and at http://www.unres.pl. Therefore, it will be only briefly outlined here. In the UNRES model^16,17,18^, a polypeptide chain is represented as a sequence of α-carbon (C^α^) atoms with virtual united peptide groups (p) located in the middle of consecutive C^α^ atoms, and united side chains (SC) are represented as ellipsoids attached to the C^α^ atoms. The force field was derived as the potential of mean force of a system of polypeptide chain(s) in solvent, where all degrees of freedom except the coordinates of the C^α^ atoms and SC centers were averaged out. The UNRES energy function is constituted by terms of local and site-site interactions as well as multibody interactions. The solvent effect is implicitly included in the SC-SC interaction function, and the force field is optimized to reproduce the structure and thermodynamics of small proteins, without being biased to the formation of amyloid fibrils. The force field used in this study was calibrated with the α-helical protein 1ENH and the β-strand protein 1E0L.

All structures of αS were extracted from REMD trajectories generated with the UNRES force field. A total of 72 trajectories were computed: 32 trajectories at the lowest temperature, 300K, and 8 trajectories at each of the following temperatures: 310K, 323K, 337K, 353K, and 370K. Each trajectory was started with two fully unfolded monomers separated by a distance of 25 Å. The conformations have been divided into three distinct classes: monomeric, dimer, and sporadic. An αS chain was classified as monomeric when each residue of one chain was at least 20Å away from the residues of the second chain. A dimeric conformation was defined as a conformation in which the two monomers had at least 10 pairs of residues in contact, with a defined contact distance of < 5.5 Å. The third class, sporadic, included all other possible conformations. The time step in the MD simulations was dt = 0.1 mtu (1 mtu = 48.9 fs is the “natural” time unit of MD^36^, which corresponds to an effective time step of 4.89 ps. Averaging of the fast degrees of freedom in UNRES results in an approximately 1,000-fold extension of the simulation timescale^37^). After discarding the first 4 million steps, 8 blocks of 10 million steps were analyzed per trajectory, with convergence assessed over the last 30 million steps (∼147 μs of effective simulation time). Only conformations at 300 and 310 K were retained for analysis, yielding ∼1 million converged structures per protein. The final dataset represents a total sampling time of ∼4.9 ms of effective simulation time. From these datasets we extract 193,024 structures containing the braid conformation using the γ/θ angle ranges defined in the Results and Discussion section. Three consecutive residues had to fall within these angle ranges to form a braid.

Moreover, 15 canonical coarse-grained MD trajectories were run at 310 K for αS using the UNRES force field, with a combined total of ∼109 million steps (∼0.53 ms of effective simulation time). The initial structure of each trajectory consisted of one fully unfolded monomer and one partially folded monomer containing a preformed braid structure, separated by a distance of at least 20 Å. After discarding the first 7.5 million steps, ∼101 million steps (∼0.50 ms of effective simulation time) were analyzed.

### Detection of α-Helices and β-Sheets

The secondary structures of α-synuclein conformations were analyzed directly from the C^α^ coordinates by using the CUTABI (CUrvature and Torsion based of Alpha-helix and Beta-sheet Identification) algorithm recently developed in our group^29^. The program is based on the analysis of 14,652 experimental structures containing less than 40% sequence identity extracted from the ASTRAL database^38,39^. The CUTABI parameters were adjusted to replicate the results obtained by the commonly used DSSP (Dictionary of Secondary Structure of Proteins)^40^ algorithm for ASTRAL database. A 93% agreement was obtained between CUTABI and DSSP with 10 – 30 times less computational cost^29^. CUTABI can only detect helices with at least 4 residues, therefore, short 3_10_ helices (comprising 3 residues) cannot be recognized by the program. The minimal size of a β-strand is set to 2 residues, i.e., a β-sheet cannot be smaller than 4 residues. Therefore, the β-bridges formed by the β-strands of 1 residue are not considered^29^.

### Coarse-Grained Angles

Assuming a constant virtual bond length between C^α^ atoms of successive residues, the conformation of a protein chain composed of *N* amino acids can be fully described using *N − 3* virtual-bond-torsional angles (γₙ) and *N − 2* virtual-bond angles (θₙ). The virtual-bond-torsional angles γₙ are defined by the positions of four consecutive C^α^ atoms: 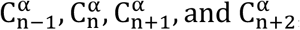, for *n* ranging from 2 to *N − 2*. The virtual-bond angles θₙ are defined by the positions of three consecutive C^α^ atoms: 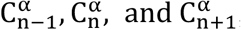, for *n* ranging from 2 to *N − 1*. These angles have clear geometrical interpretations: the virtual-bond angles θₙ correspond to the discrete local curvature, while the virtual-bond-torsional angles γₙ represent the local torsion of the chain formed by the successive virtual C^α^−C^α^ bonds^17,21,41^. Mathematically, local curvature and torsion provide a complete set of local order parameters, fully describing the geometry of a space curve and serving as a fundamental framework for understanding protein folding^41^.

### Ramachandran Angles

Assuming ideal bond lengths and angles within the peptide backbone, the three-dimensional conformation of a protein can also be fully described by the sequence of its backbone dihedral angles φ and ψ. For each amino acid residue (except at the termini), the φ angle is defined by the torsion around the N–C^α^ bond and involves the atoms 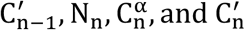. The ψ angle is defined by the torsion around the C^α^–C′ bond and involves the atoms 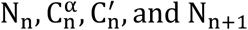. These angles capture the rotational degrees of freedom of the protein backbone and reflect the local conformational state of each residue. In the Ramachandran plot^42^, φ and ψ values are used to identify sterically allowed regions of conformational space, highlighting preferred secondary structure motifs such as α-helices and β-sheets. From a mathematical perspective, the φ/ψ angle pair for each residue serves as a minimal set of local coordinates to describe the protein’s backbone geometry and plays a central role in understanding protein structure and folding.

## Results and Discussion

### Structures Adopted by the 35-55 Fragment of α-Synuclein

Apart from the work by Yu *et al*.^31^, a recent experimental studies showed that this region, named ‘master controllers’ regions, P1 (residues 36-42) and P2 (residues 45-57), regulates aggregation of αS and can be considered as a target for future therapeutic intervention^43,44^. Moreover, in our recent study^45^, by analyzing the experimental structures of the micelle-bound, tetrameric, and fibrillar αS in terms of a kink (heteroclinic standing wave solution)^46^ of a generalized discrete nonlinear Schrödinger equation, we illustrated that this region plays an important role in the stability of αS structure. Therefore, we carried out a detailed analysis of the 35-55 region using all isolated monomers extracted from our coarse-grained REMD simulations, which are 165 times longer than simulations by Yu *et al*.^31^. The goal of our analysis was not only to compare our findings with previously published results, but also to determine whether this region can adopt conformations other than b-hairpin.

Secondary structures such as α-helices or β-sheets can be easily identified from the bond (θ) and dihedral (γ) angles between the virtual bonds joining consecutive α-carbon (C^α^) atoms by using the CUTABI algorithm^29^. Here, we aim not only to identify helices or sheets but also to detect random coils and other structural motifs. To this end, we computed the number of intra-chain contacts within residues 35–55 using a cutoff distance of 8 Å excluding contacts with immediate neighboring residues (*i*+1 and *i*+2). The probability distribution of contacts at this cutoff clearly reveals four distinct clusters (Figure 2). Notably, among several tested cutoff distances, 8 Å provided the best separation between these four conformational states. To visualize the structural content of these clusters, we selected the local maxima of the contact-number distributions (peak maxima) and computed contact maps for all structures exhibiting the corresponding number of contacts within the 35–55 region. The first cluster, characterized by the lowest contact count (2 contacts), corresponds to a random-coil ensemble lacking well-defined secondary structure within the 35–55 residue segment. The second cluster, with 26 contacts, predominantly adopts a hairpin conformation. The third cluster, with 35 contacts, also forms a hairpin, but with the turn positioned centrally within the 35–55 window. In contrast, for the second cluster the hairpin turn is shifted toward lower residue indices, such that the N- and C-terminal ends of the strands fall partially outside the 35–55 region and are therefore not counted as contacts, explaining the lower contact number.

**Figure 2.**
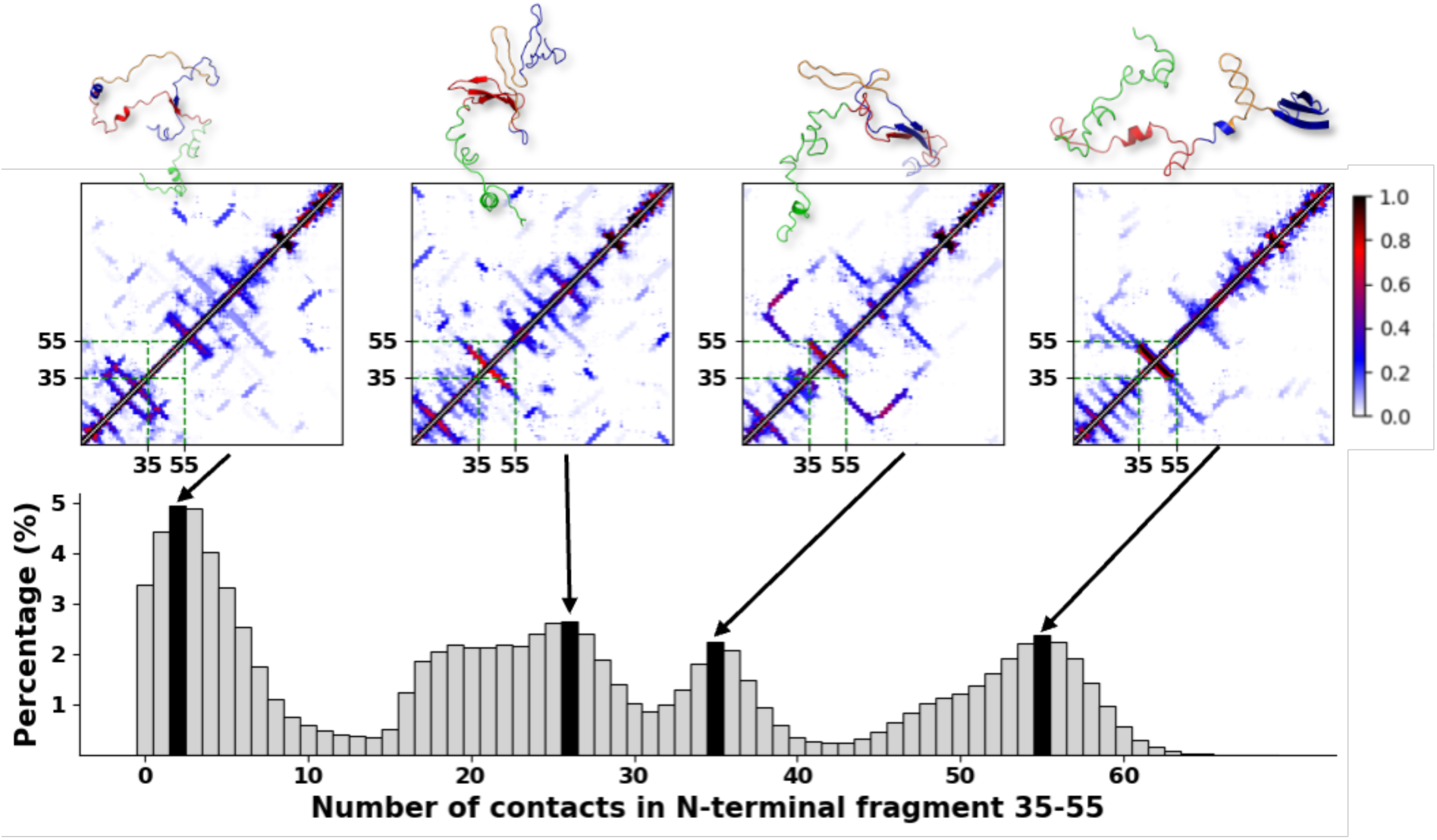
Percentage of contact probabilities within an 8 Å cutoff for residues 35–55 across all monomeric WT αS structures. For each black bar in the histogram, the corresponding 2D contact map shown above represents the ensemble of structures belonging to the associated cluster; each pixel indicates the probability of contact between a pair of residues according to the color scale. Representative 3D structures of each cluster are displayed above the 2D maps, with residues 1–61 shown in blue, the 35–55 region highlighted in orange, residues 62–95 (NAC region) in red, and residues 96–140 (C-terminal region) in green.

The most aggregated cluster, characterized by a very high number of contacts (55 contacts), exhibits a contact map containing a broad diagonal band reminiscent of a hairpin, although significantly wider, indicating the presence of a dense and extended network of contacts. This cluster predominantly, though not exclusively, contains a novel structural motif that we termed a *braid*. Inspection of the three-dimensional conformations associated with this motif reveals a hairpin-like fold twisted into a helical shape. The structural and dynamical properties of the braid motif are described in the following section.

### Braid Type Structures

Based on the histogram panel in Figure 2, the presence of the braid and braid-like structures in the datasets is somewhat significant, consequently it may have a crucial impact not only on possible folding of the rest of the chain in this region, but also on aggregation with another monomer of αS. Since the braid structure basin is quite wide, 43-63 contacts, it contains many not-fully formed braids but also other type of strongly aggregated structures; therefore, we selected the most perfectly formed braid structure as a reference, and computed RMSD from the reference structure. It appeared that only structures with the RMSDs less than 3.5 Å keep fully-formed braid conformations, which is 3.1% of entire dataset. The results presented herein are based on perfectly formed braid structures.

To better characterize the geometry of the braid, we analyzed several structural parameters, including angular- and contact-based descriptors. We first examined the coarse-grained dihedral angles γ and bond angles θ. Heatmap of the two-dimensional probability density function (PDF) of γ and θ angles pair is shown in Figure 3, together with the corresponding probability histogram of residues located within the region highlighted by a square on the heatmap. Two main regions, referred to as the “*body*” and the “*turn*” regions, as shown in Figure 3. The braid is formed primarily between residues 37 and 51, with γ and θ angles of most of its residues ranged from 0° to 80° and from 105° to 130°, respectively, for the “*body*”, and with γ and θ angles ranged from –80° to 0° and from 60° to 110°, respectively, associated with the “*turn*”. The turn is primarily formed by residues 43 and 44, while residues 42 and 45 serve as transition points between the turn and the body of the structure (Figure S1).

**Figure 3.**
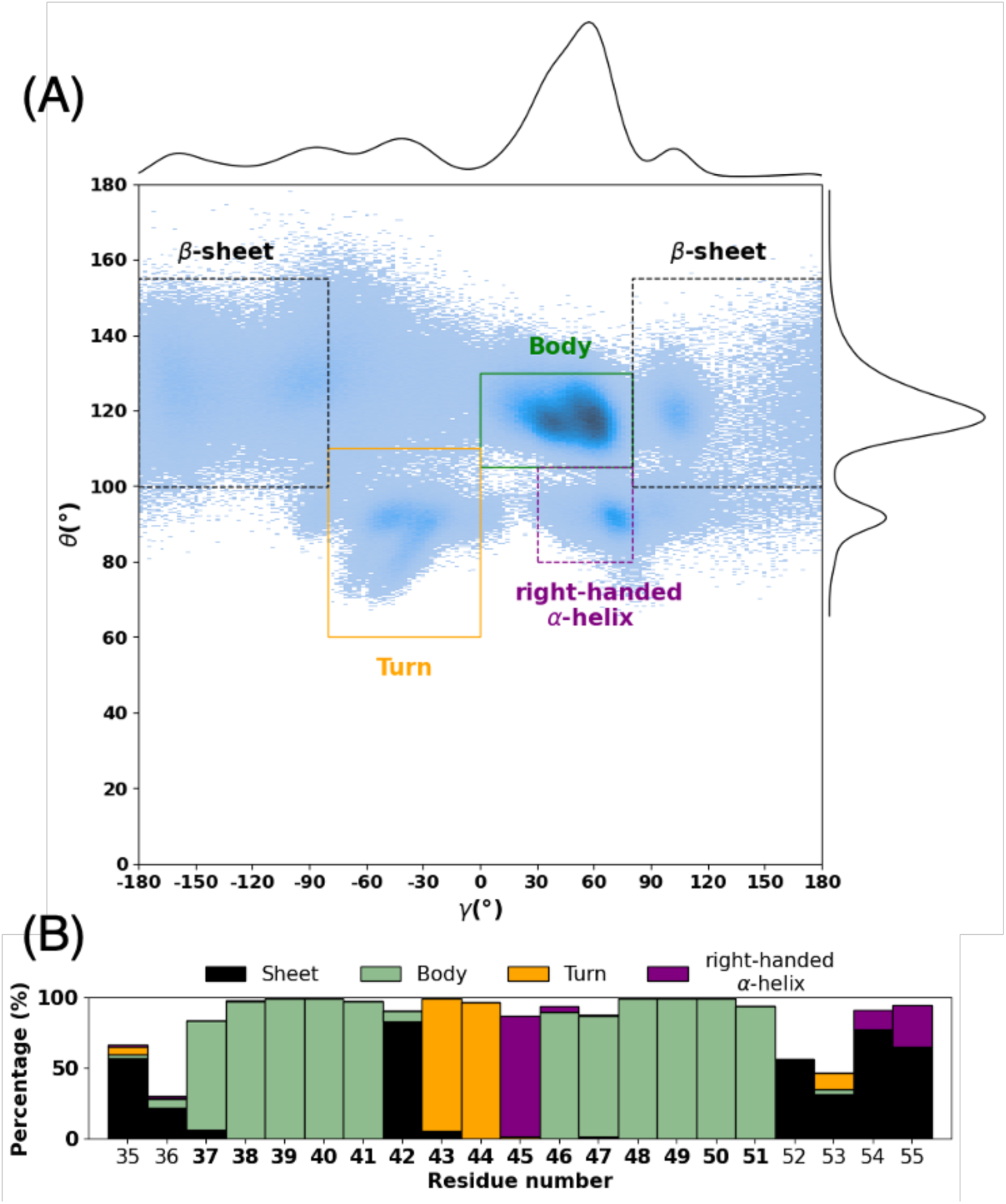
(A) Heatmap of the 2-D PDF of γ and θ angles for datasets containing braids. Black dashed rectangles indicate the β-sheet angle ranges, while purple dashed rectangles correspond to the right-handed α-helix as defined by CUTABI. The *turn* and *body* rectangles represent newly defined angle ranges, detailed in the main text. (B) Stacked bar plot showing the probability of each conformational state, based on the angle ranges defined in panel A, for residues 35 to 55.

As was mentioned above, the *body* region lies between the β-sheet and right-handed α-helix as defined by the CUTABI software^29^. This suggests that the braid is more extended than a typical right-handed α-helix, characterized by a larger θ angles, but less planar than a β-sheet, as indicated by a smaller γ angles. Moreover, the regions of θ and γ angles associated with the *turn* correspond the regions of left-handed α-helices.

To construct a heatmap of the two-dimensional PDF of φ and ψ angles, the dataset containing braid conformations in coarse-grained coordinates were converted to an all-atom representation using Multiscale Modeling Tools for Structural Biology (MMTSB)^47^. As expected, the two-dimensional map of the ϕ and ψ angles (Figure 4) is more complex than that of the γ and θ angles (Figure 3).

**Figure 4.**
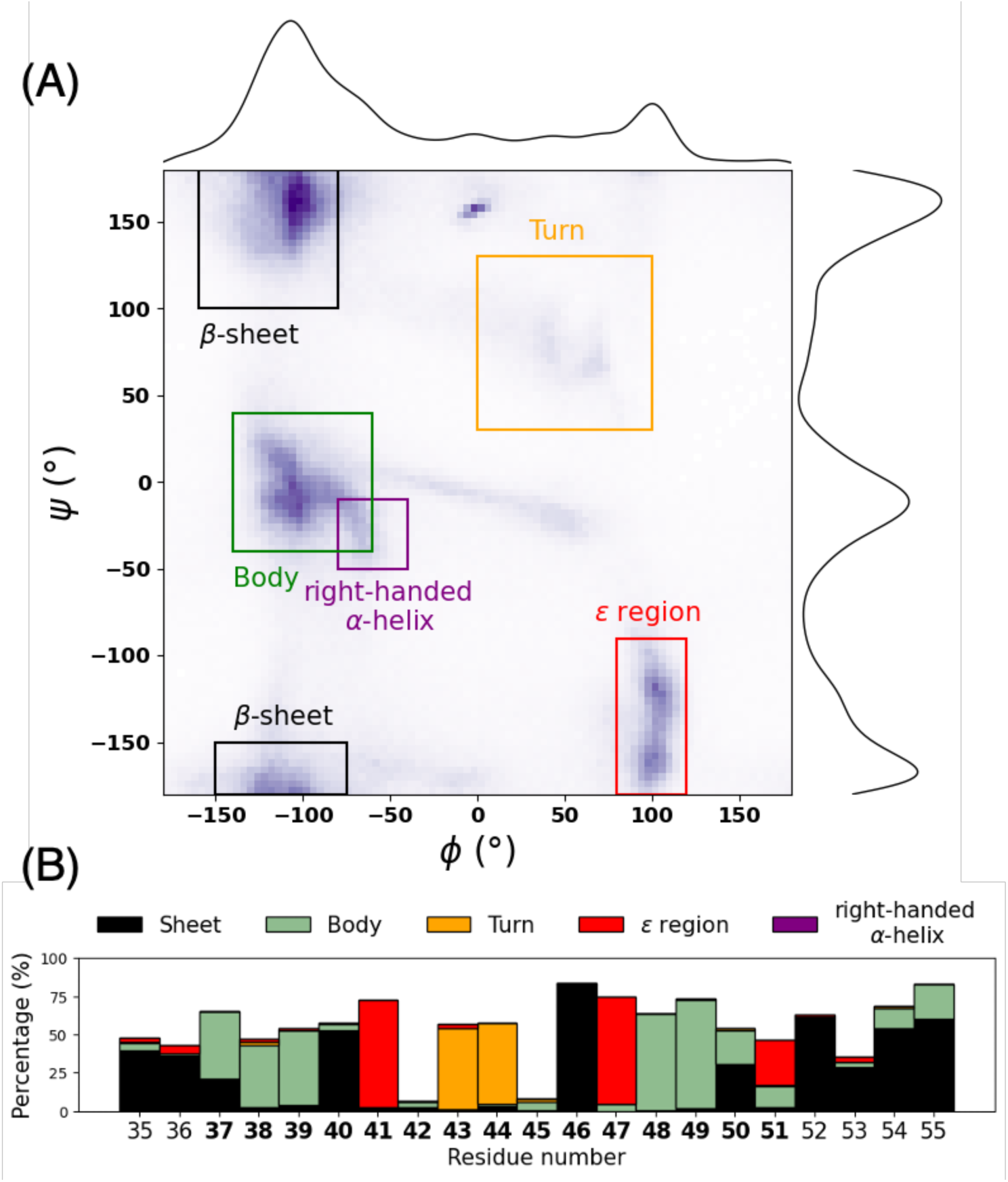
(A) The ψ/ϕ angles heatmap for braid region. Well-known α-helices regions and β-sheet region are highlithed in blacklines and purple lines, respectively. (B) Stacked bar plot of the probability of being in a *body*, *turn*, or other conformation for residues 35 to 55, based on the range defined by the panel (A).

Five regions can be clearly identified in Figure 4. First, the β-sheet region (the top-left region in quadrant II and the bottom-left region in quadrant III) corresponds mainly to residues outside the braid structure (35, 36, 52-55); however, two residues associated with the braid structure, 40 and 46, as well as three glycine residues, 41, 47 and partially 51, are also found in the β-sheet region. The second distinct region, termed the *body* region, is located between the right-handed α-helix and β-sheet regions and spans quadrants II and III. This region corresponds to residues associated with the braid structure. Notably, the *body* region coincides with a sparsely populated area, or even a void, in most traditional Ramachandran plots. Yet, the φ/ψ angles of residues within the *body* of the braid fall precisely within this underrepresented space. This observation reinforces the intermediate character of the *body*, which structurally lies between helical and sheet conformations, consistent with our previous observation based on the γ and θ angles. Compared with the right-handed α-helix, the *body* adopts slightly lower ϕ and higher ψ values, making the braid structure somewhat more extended. A region corresponding to left-handed α-helices, located in quadrant I and characterized by positive φ and ψ values, includes residues 43 and 44 of the *turn* region. These residues are largely isolated within their regions, exhibiting only slight variability (Figure S2). These results are in harmony with the above-presented PDF along the γ/θ angles shown in Figure 3. The final distinctive conformational region is the ε region, located in the sparsely populated lower portion of the left-handed quadrant IV of the Ramachandran plot. This region reflects a relatively extended backbone conformations and is populated predominantly by glycine residues^48^ (Figure S2). Glycine is uniquely capable of occupying the ε region because its lack of a β-carbon (C^β^) minimizes steric hindrance, allowing access to backbone torsion angles with positive φ and negative ψ values that are energetically unfavorable for most other amino acids. As a result, glycine can readily sample conformations associated with left-handed helical geometry and other unusual backbone states. Transient sampling of these extended dihedral-angle conformations within residues 35–55 may facilitate exposure of otherwise buried hydrophobic surfaces, potentially increasing the likelihood of aberrant intermolecular interactions and oligomer formation^49,50^.

The braid can also be characterized in terms of residue contacts. As shown in the contact map in Figure 5, residues in the ranges 36–43 and 47–53 consistently form at least two contacts even when a stringent cutoff of 4.5 Å is applied, highlighting the compactness of this structure.

**Figure 5.**
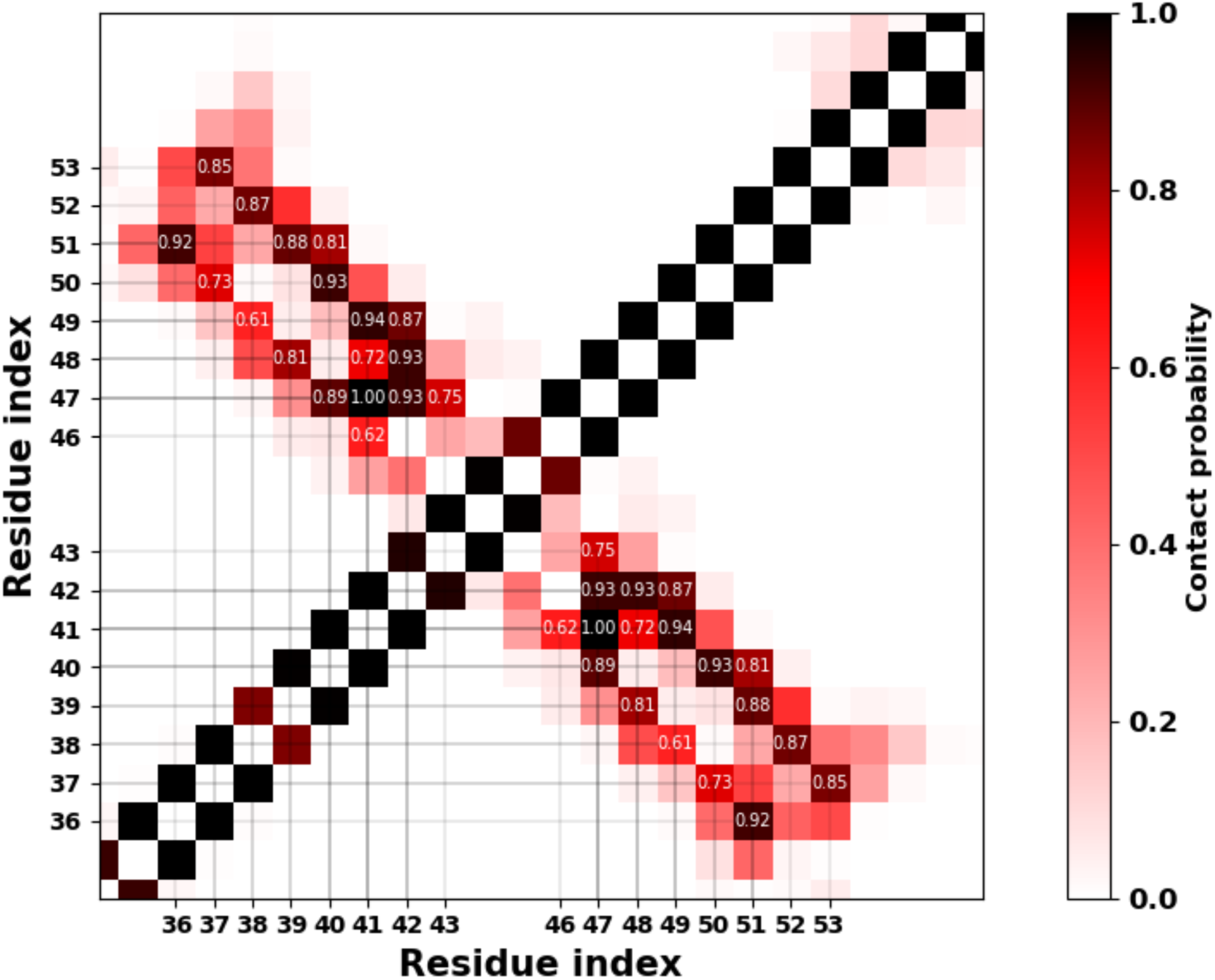
Contact map of the braid region with a 4.5 Å cutoff. Probability is written for a pair of residues if the probability is higher than 0.6. Corresponding residues are given in X and Y axis.

The braid is distinctive due to its unique geometry and, above all, its compactness. This type of structure packs a large number of residues within a very small surface area. It encompasses 15 residues. Consequently, the steric hindrance is significant. This compactness is facilitated by a high proportion of hydrophobic residues, five valines, one alanine, and one leucine, representing nearly half of the total residues, as well as three glycines, which lack side chains and thus reduce steric bulk.

### Steps of Braid Formation

To better understand the formation of the braid structure, we need to investigate further how it is formed and if it depends on the α-synuclein environement or on its intrinsic sequence. To reduce the cost of computation we performed 40 canonical UNRES MD trajectories of a short 35-55 segment of α-synuclein at 300 K. For each trajectory, 50 000 steps (i.e. ∼ 0.25 ns formal time and effectively ∼ 0.25 μs) were run, starting from the fully extended conformations.

We first analyzed the 35–55 segment and computed the RMSD for each trajectory using a perfect braid structure extracted from our dataset as a reference. As illustrated in Figure 6, many trajectories rapidly reach low RMSD values, indicating that braid formation occurs early, although at different simulation steps. The PDF, shown in black on the right side of the plot, reveals seven local maxima corresponding to distinct conformational states. State 1, which has the lowest RMSD and is closest to zero, exhibits a clear braid geometry, although it displays noticeable instability, fluctuating between states 1 and 2. State 2 shows a small deviation localized in the turn of the braid, suggesting that this region is flexible. State 3 is a rare state characterized by a larger deformation of the turn. State 4 is structurally close to state 1; however, a deviation of the central axis of the braid leads to a significantly higher RMSD. States 5 and 6 occur at similar RMSD values and correspond to transitional braid conformations, indicating that at least two folding mechanisms can lead to braid formation. In state 5, folding is initiated at the turn, whereas in state 6 it starts at the base of the braid. Both states are highly noisy and unstable, preceding the transition to lower-RMSD states that are relatively stable. In terms of probability, states 2 and 4 are more populated than the other intermediate states. Finally, although two trajectories (light green and pink) fail to fold into a braided conformation, the majority successfully reach the braid structure. The representative structure of the highest-RMSD state (state 7) does not exhibit any braid-like features and is therefore not shown in the Figure 6.

**Figure 6.**
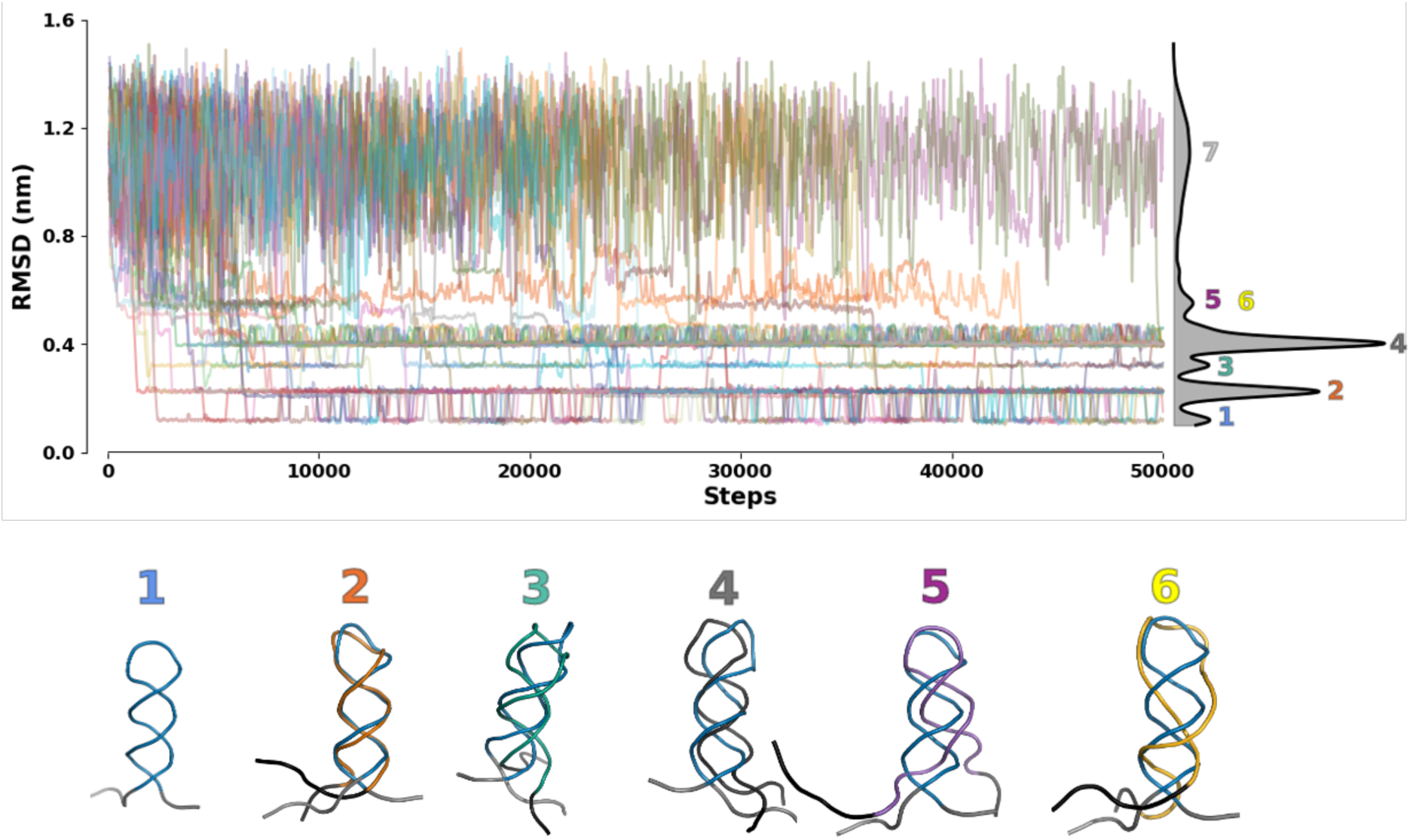
Root-mean-square deviations (RMSDs) from the reference braid structure (perfect braid) as functions of simulation steps for 40 individual MD trajectories, generated with the UNRES force field, starting from the fully extended conformation of segment 35–55 corresponding to the braid region. A sliding-window average was applied to smooth the curves and improve readability. The global PDF is shown alongside the trajectories. Three-dimensional representations of the structures corresponding to each local maximum are provided using the same numbering and color code. For each structure, state 1 is retained to allow direct comparison with the reference braid conformation.

These results indicate that 35-55 segment of αS is intrinsically capable of forming a braid structure on its own. However, in full-length α-synuclein, this conformation is observed in only 3.1 % of the ensemble, suggesting that the surrounding protein environment imposes constraints that limit the formation or stability of the braid in this segment.

### All-Atom Simulations

To assess the dependence of braid stability on the force field, we examined its stability using all-atom simulations. Due to the large difference in timescales between all-atom and coarse-grained force fields, it is not feasible to simulate the folding of the braid starting from an extended structure, as can be done with UNRES, a force field that has been successfully used to simulate *ab initio* protein folding. Therefore, we adopted the following strategy. First, we randomly selected ten braid structures from our datasets and used them as initial conformations for ten 40 ns trajectories performed with two different all-atom force-fieds, CHARMM36 (C36)^51^ and AMBER^52^, at two temperatures (250K and 300K). The RMSD of each trajectory as a function of time is shown in Figure 7, together with RMSD distributions calculated for the intervals from 0 ns to 5 ns (black) and from 5 ns to 40 ns (orange).

**Figure 7.**
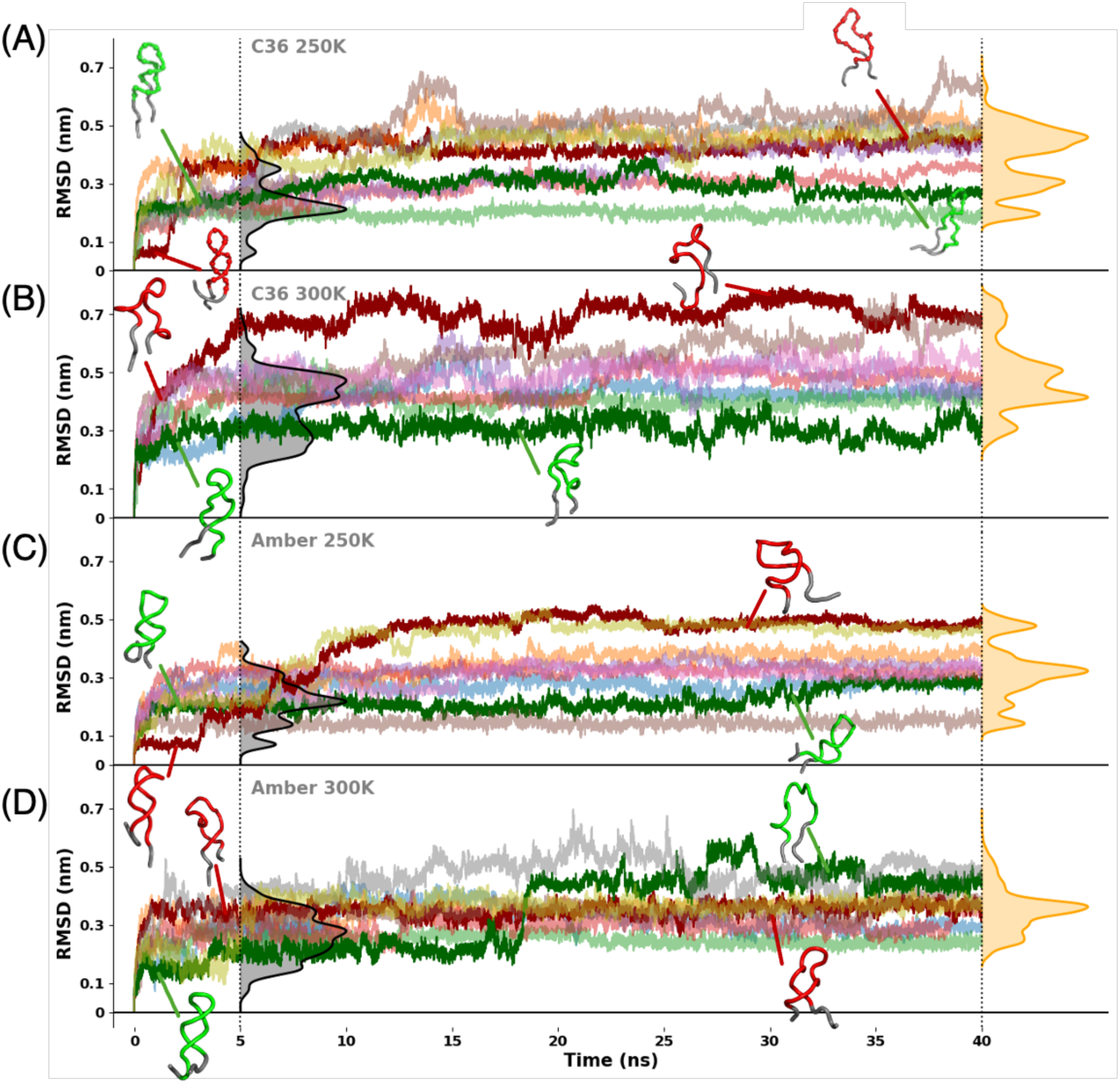
RMSDs from the braid structure as functions of time for 10 trajectories, generated with the CHARMM36 (C36) (panels A and B) and AMBER (panels C and D) force fields at 250 K and 300 K temperatures, starting from the braid structures. RMSD distributions for the first 5 ns are shown in black, and distributions from 5 ns to 40 ns are shown in orange. Selected trajectories are highlighted in color to illustrate representative three-dimensional conformations.

In the trajectories generated with the CHARMM36 force field at 250 K (panel A), the RMSDs exhibit transitions between three distinct states within the first 5 ns. A first, very low-RMSD state at ∼0.06 nm is observed in only one trajectory (maroon). Its three-dimensional representation shows a nearly perfect braid; however, after ∼2 ns this structure rapidly unfolds to ∼0.35 nm, converging with the other trajectories, which reach this state almost immediately after the start of the simulation. A second intermediate state (state 2) at ∼0.22 nm is the most populated and corresponds to a braid structure displaying moderate deformations. After 5 ns, most trajectories transition to a higher-RMSD state at ∼0.50 nm, corresponding to an unfolded braid adopting a large-loop conformation. Only three trajectories remain at lower RMSD values (∼0.20 nm and ∼0.30 nm), retaining partial braid features. In particular, the green trajectory preserves a single loop near the turn region.

At 300 K (panel B), a more physiological temperature, the RMSD increases more rapidly in most trajectories, reaching ∼0.47 nm. Only two trajectories (blue and green) remain at lower RMSD values (∼0.27 nm) for an extended time. The blue trajectory eventually converges toward higher RMSD values, whereas the green trajectory remains stable at low RMSD. Three-dimensional inspection of the latter reveals retention of a loop close to the braid turn. After 5 ns, most trajectories either remain in the same unfolded state or transition to even higher RMSD values (maroon and light brown trajectories), corresponding to fully unfolded conformations. In contrast, the green trajectory, which remains at lower RMSD, preserves a loop near the turn that closely resembles the initial structure.

Simulations using the AMBER force field show behavior similar to CHARMM36 at 250 K (panel C), including one trajectory that maintains a stable braid-like structure for approximately 3 ns. Most trajectories populate two states at ∼0.14 nm and ∼0.22 nm, both of which retain two loop features. Ultimately, most trajectories converge to an RMSD of ∼0.30 nm, except for the light brown trajectory, which remains stable at ∼0.16 nm throughout the simulation, and two others (maroon and yellow) that transition to a higher RMSD (∼0.50 nm), corresponding to a single-loop unfolded conformation.

At 300 K (panel D), trajectories generated with the AMBER force field display behavior comparable to those observed at 250 K, with an initial state around ∼0.2 nm before 5 ns and a dominant state at ∼0.35 nm afterward.

Overall, these results demonstrate that the braid structure can form and persist transiently in all-atom force fields, both in AMBER and CHARMM36, despite its compactness. However, this state is not fully stable for these all-atom Hamiltonian parametrizations and rapidly unfolds, indicating that the braid corresponds to a metastable conformational state in these force fields. The spontaneous folding of the braid in all-atom simulations is expected to have a low probability. Therefore, only the UNRES data for the braid structures are discussed in the remainder of the paper.

### Impact of the braid on the global structure of αS

To understand the general behavior of the braid structure and its impact on the rest of the protein, we constructed the contact maps (CMs) of all braid conformations observed in WT αS (Figure 8, left panel) compared to the CMs of all other structures found in our datasets (Figure 8, right panel).

**Figure 8.**
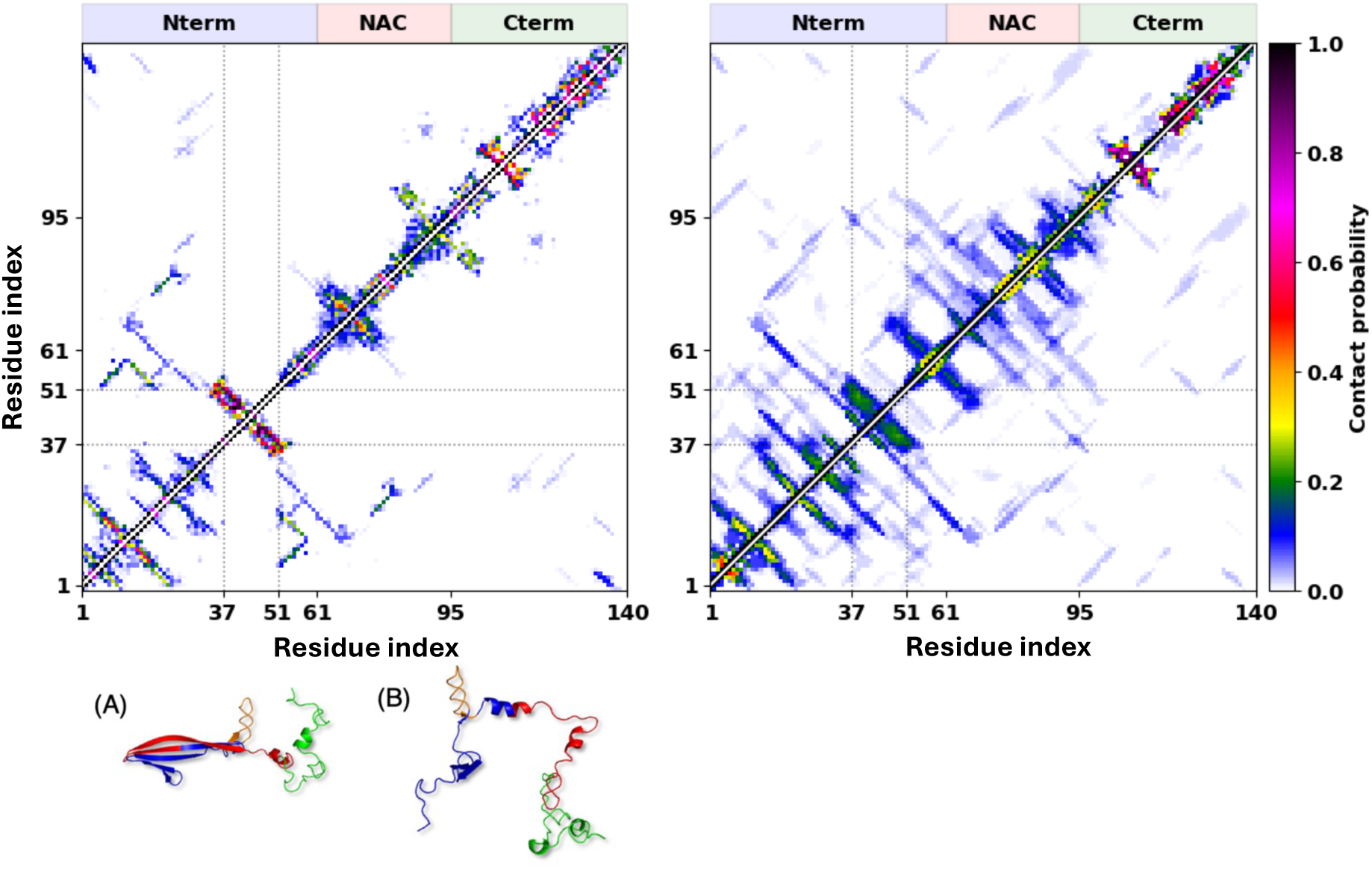
Contact maps (5.5 Å cutoff) of WT αS conformations containing the braid motif (left panel), together with representative structures (panels A and B), and of all other conformations in the dataset (right panel). The rectangles outlined by dotted grey lines correspond to the 37-51 residue segment. The different protein regions are indicated above the contact maps.

In the CMs shown in Figure 8, the braid is easily identifiable as a thick trace perpendicular to the diagonal, reflecting the numerous residue-residue contacts formed within the structure under the selected cut-off radius. The CM indicates that the braid does not interact with the remainder of the protein, unlike all other structures of the 37-51 segment, including the second representative structure - the β-hairpin^53^. The representative structures illustrated in Figure 8 reveal that, in the presence of the braid, the NAC and N-terminal regions of WT αS can adopt markedly different organizations. In one case [structure (A)], these regions assemble into an extended β-sheet composed of multiple strands arranged within the same β-plane. In contrast, structure (B) shows the two regions remaining largely separated, accompanied by the formation of an α-helix at their interface. These observations highlight the structural heterogeneity associated with braid-containing conformations and suggest that the braid can coexist with distinct long-range arrangements of the protein.

Another interesting observation is that long-range contacts are absent not only between the braid region and the remainder of the protein but are also greatly reduced between other parts of αS. In other words, the braid structure appears to hinder the formation of long-range intramolecular interactions, leaving the N-terminus, NAC region, and C-terminus solvent-exposed and potentially facilitating fibril nucleation.

We also investigated the formation of other secondary-structure elements in α-synuclein in the presence or absence of the braid. Figure 9 shows that α-helical content within the NAC region increases when the braid is formed. Conversely, β-sheet formation decreases both in the vicinity of the braid and within the NAC region, likely as a consequence of the increased helix formation. In contrast, the very beginning of the N-terminal region (residues <30) is only minimally affected by the presence of the braid, while the C-terminal region appears to remain completely unaffected.

**Figure 9.**
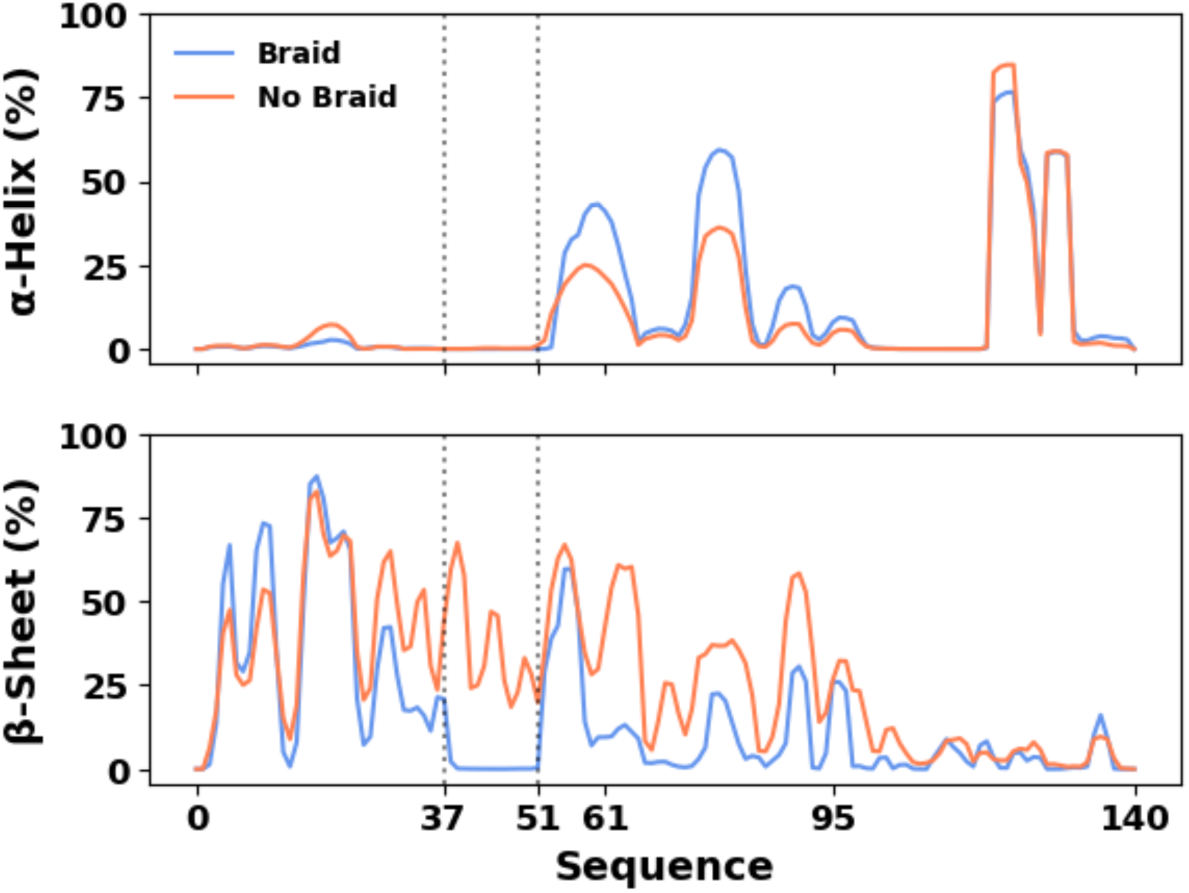
Top panel: probability (in %) for each residue in αS to form an α-helical conformation in the presence (blue) or absence (orange) of the braid. Bottom panel: same as the top panel, but showing the probablity of β-seet formation for each residue.

### Braid structures in dimers

In order to determine whether the braid structures observed in monomers persist upon dimerization, we extended our analysis of the braid structure to dimers. As shown in Figure 10, the inter-chain contact maps for dimers with (left panel) and without (right panel) a braid (i.e., at least one monomer must adopt a braid conformation for the dimer to be classified as ‘with braid’) indicate that contacts involving the braid region are substantially reduced or entirely absent in dimers with a braid compared with those without a braid. The only region that forms strong intermolecular contacts with the braid region corresponds to the beginning of the N-terminus. In contrast, in dimers consisting of monomers without braid structures, intermolecular contacts are observed between different regions of the monomers, including fibril-like contacts within the NAC region (right panel of Figure 10). Owing to the limitations of the present REMD-based approach, it remains unclear at what stage the braid forms. However, our results suggest that the braid most likely forms prior to dimerization. The results illustrated in right panel are consistent with previous studies indicating that the NAC region forms the core of αS fibrils.

**Figure 10.**
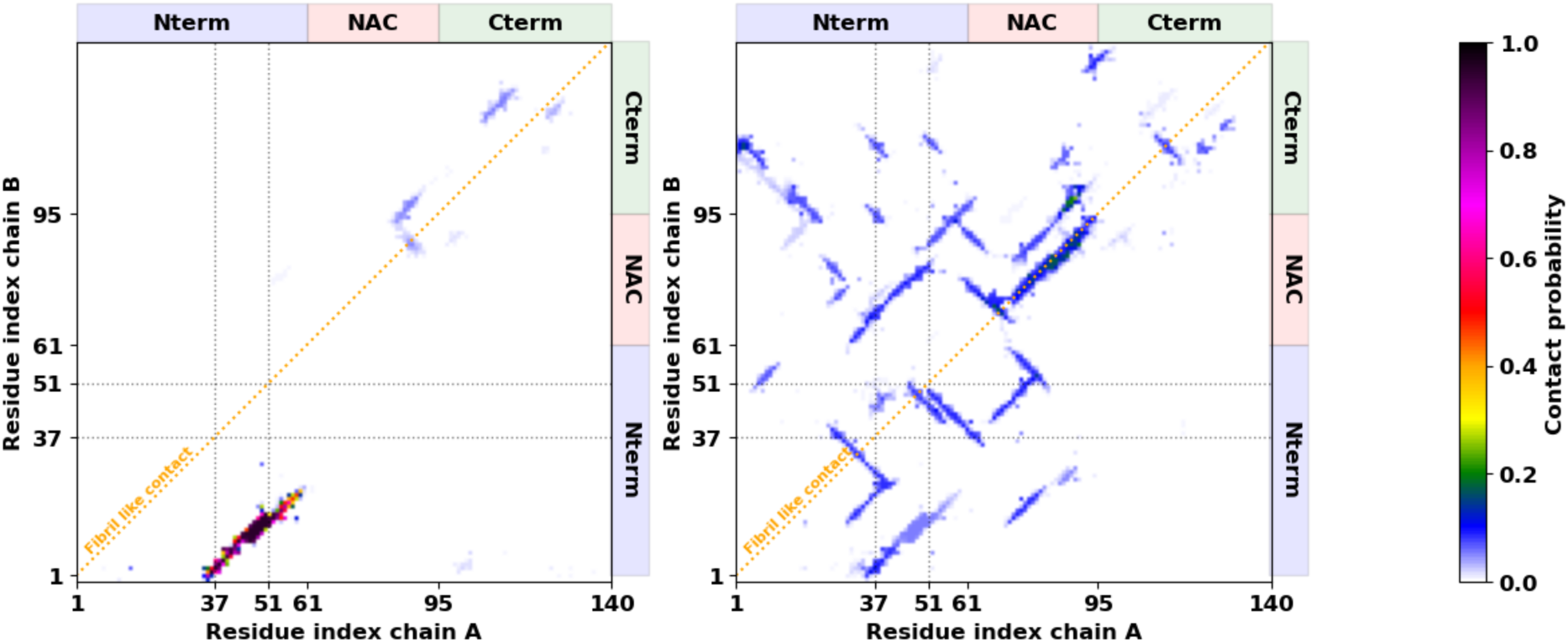
Inter-chain contact maps calculated using a 5.5 Å distance cutoff for α-synuclein dimer datasets. The left panel shows dimers in which at least one monomer contains a braid structure, whereas the right panel shows dimers in which neither monomer contains a braid. The rectangles outlined by dotted grey lines correspond to the 37-51 residue segment. The different protein regions are indicated above and on the right side of the contact maps. The orange dashed lines represent the diagonals corresponding to fibril-like contacts, where identical residues from both chains are in contact.

Moreover, it is of interest to investigate how the braid conformation can persist within dimeric assemblies and hinder inter-chain folding and self-association. To address this question, we performed canonical MD simulations of αS, in which the initial structure consisted of one fully unfolded monomer (chain B) and one partially folded monomer containing a preformed braid structure (chain A), separated by a distance of at least 20 Å. In particular, we ran 15 trajectories at 310K, with a total effective simulation time of ∼ 0.5 ms. By computing the RMSDs of the braid structure as a function of time (not shown), we monitored its stability and found that the braid structure remained intact throughout all trajectories. The inter-chain contact maps and histograms of contacts between the regions, plotted in Figure 11, indicate that the braid regions either do not interact (using a 5.5 Å cutoff distance) or interact only minimally (using a 8.0 Å cutoff distance). In addition, the braid structure formed in chain A interacts with all regions of the chain B, especially with the C-terminus.

**Figure 11.**
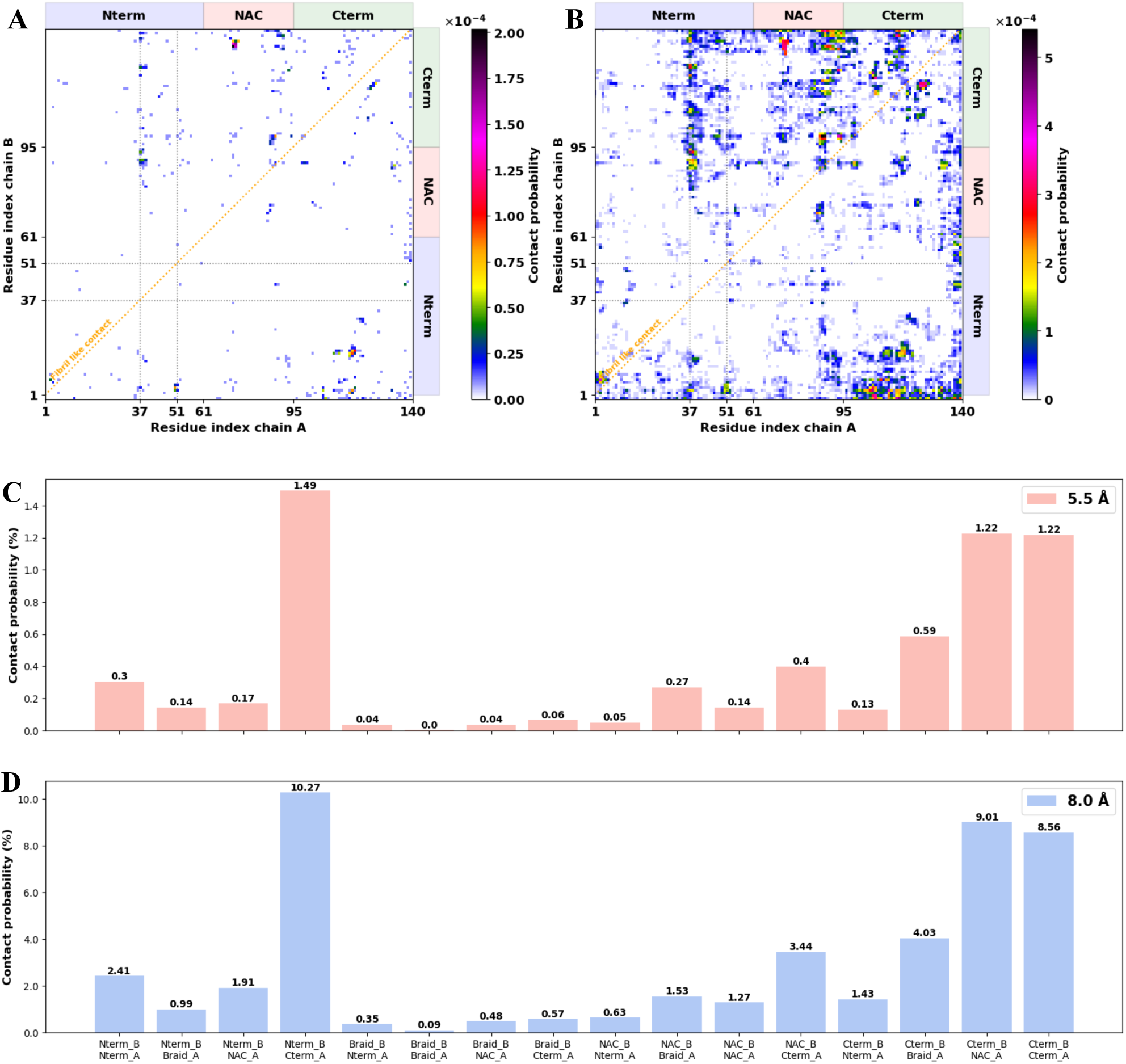
Inter-chain contact maps calculated using 5.5 Å (panel A) and 8.0 Å (panel B) distance cutoffs for α-synuclein datasets of sporadic interactions, and histograms of inter-chain contact probabilities between protein regions calculated using 5.5 Å (panel C) and 8.0 Å (panel D) distance cutoffs. In panels A and B, the rectangles outlined by dotted grey lines correspond to the 37-51 residue segment. The different protein regions are indicated above and on the right side of the contact maps. The orange dashed lines represent the diagonals corresponding to fibril-like contacts, where identical residues from both chains are in contact.

Interestingly, the presence of the braid structure appears to impede interactions between the NAC regions, including fibril-like contacts. In general, the regions surrounding the braid can be characterized as being largely devoid of inter-chain contacts. Frequent interactions are observed between the N-terminus of chain B and the C-terminus of chain A, between the NAC region of chain A and the C-terminus of chain B, and between the C-termini of chains A and B. However, very few fibril-like contacts are observed among these interactions.

These findings are consistent with previous studies indicating that the so-called ‘master controllers’ regions, P1 (residues 36-42) and P2 (residues 45-57), which overlap with the braid region, regulate aggregation of αS^43,44,54^. In other words, the conformations adopted by the ‘master controllers’ regions may modulate aggregation process. Overall, despite the sporadic nature and low probability of these contacts, the present results reveal a characteristic pattern of interactions involving this region.

It should be noted that, in our recent study of αS dimers^30^, we found that fibril-like contacts within the NAC region are significantly more abundant in WT than in the E46K and A53T mutants. Conversely, the number of fibril-like contacts within the N-terminal region is significantly higher in the E46K and A53T mutants than in WT. These results suggest that even a single amino acid substitution within the ‘master controllers’ regions can modulate the aggregation propensity of αS.

## Conclusions

Three distinct conformations – the β-hairpin, braid, and random coil – were identified in a short fragment encompassing residues 37-51 of WT αS and its familial mutants A30P, A53T, and E46K. In this study, we characterized the braid, a previously unidentified secondary structure observed in WT αS, using the UNRES force field. Although we could not definitively rule out the possibility that the UNRES force field overestimates the stability of the braid, two all-atom simulations demonstrated that this structure is sterically feasible and metastable. The 37-51 fragment, in which the braid structure forms, lies within the P1 and P2 regions, which are known to regulate α-synuclein aggregation. We have shown that the braid structure appears to hinder the formation of long-range intramolecular interactions, leaving the N-terminus, NAC region, and C-terminus solvent-exposed and potentially facilitating fibril nucleation. Interestingly, in dimers, the braid structure appears to impede intermolecular contacts, especially homotypic intermolecular interactions, thereby slowing or preventing aggregation. Upon unfolding of the braid structure, however, intermolecular contacts, including homotypic interactions, increase noticeably, indicating the initiation of an aggregation process. These results suggest that the braid structure may form prior to dimerization. Furthermore, our results suggest that the 37-51 segment can adopt extended backbone conformations characterized by unusual dihedral-angle values. Sampling of these conformations may facilitate the exposure of otherwise buried hydrophobic surfaces, thereby increasing the likelihood of aberrant intermolecular interactions and the formation of partially folded intermediates. Such intermediates may subsequently assemble into neurotoxic oligomeric species. Notably, recent studies^55^ have suggested that amyloid-β (Aβ), one of the main players in Alzheimer’s disease, may also adopt a braid-like conformation. The observation of similar structural motifs in two aggregation-prone proteins associated with major neurodegenerative disorders raises the possibility that braid conformations may represent a broader structural feature relevant to protein misfolding and amyloid formation.

Taken together, these findings suggest that the braid structure may play an important role in the aggregation behavior and pathogenic mechanisms of both αS and Aβ. However, the biological significance of this conformation remains to be established. Therefore, experimental validation of the existence of the braid structure and its potential role in protein aggregation is planned for future studies.

## Acknowledgements

The simulations were performed using HPC resources from DSI-CCuB (Universite Bourgogne Europe). The work is part of the project SEPIA supported by the EIPHI Graduate School (contract ANR-17-EURE-0002), the Conseil Regional de Bourgogne-Franche-Comte, and the European Union through the PO FEDER-FSE Bourgogne 2021/2027 program.

## Author Contributions

**Adrien Guzzo:** Data curation (equal); Formal analysis (equal); Investigation (equal); Methodology (equal); Software (equal); Visualization (equal); Writing – original draft (equal); Writing – review and editing (equal). **Patrice Delarue:** Data curation (equal); Formal analysis (equal); Investigation (equal); Methodology (equal); Software (equal); Visualization (equal). **Khatuna Kachlishvili:** Data curation (equal); Formal analysis (equal). **Adrien Nicolaï:** Data curation (equal); Formal analysis (equal). **Patrick Senet:** Conceptualization (lead); Data curation (equal); Formal analysis (equal); Funding acquisition (lead); Project administration (lead); Resources (lead); Supervision (lead); Writing – original draft (equal); Writing – review and editing (equal). **Gia G. Maisuradze:** Conceptualization (lead); Data curation (equal); Formal analysis (equal); Supervision (lead); Writing – original draft (lead); Writing – review and editing (lead).

## Data Availability

The data that support the findings of this study are available from the corresponding authors upon reasonable request.

## References

1 Spillantini, M. G.; Schmidt, M. L.; Lee, V. M.-Y.; Trojanowski, J. Q.; Jakes, R.; Goedert, M. α-synuclein in Lewy bodies. Nature 1997, 388, 839–840. 10.1038/42166

2 Burré, J.; Vivona, S.; Diao, J.; Sharma, M.; Brunger, A. T.; Südhof, T. C. Properties of native brain α-synuclein. Nature 2013, 498, E4–E6. 10.1038/nature12125

3 Theillet, F. X.; Binolfi, A.; Bekei, B.; Martorana, A.; Rose, H. M.; Stuiver, M.; Verzini, S.; Lorenz, D.; van Rossum, M.; Goldfarb, D.;, et al. Structural disorder of monomeric α-synuclein persists in mammalian cells. Nature 2016, 530, 45–50. 10.1038/nature16531

4 Bartels, T.; Choi, J. G.; Selkoe, D. J. α-synuclein occurs physiologically as a helically folded tetramer that resists aggregation. Nature 2011, 477, 107–110. 10.1038/nature10324

5 Wang, W.; Perovic, I.; Chittuluru, J.; Kaganovich, A.; Nguyen, L. T. T.; Liao, J.; Auclair, J. R.; Johnson, D.; Landeru, A.; Simorellis, A. K.;, et al. A Soluble α-synuclein construct forms a dynamic tetramer. Proc. Natl. Acad. Sci. U.S.A. 2011, 108, 17797–17802. 10.1073/pnas.1113260108

6 Fink, A. L. The aggregation and fibrillation of α-synuclein. Acc. Chem. Res. 2006, 39, 628−634. 10.1021/ar050073t

7 Goldberg, S. S.; Lansbury, P. T. Jr. Is there a cause-and-effect relationship between α-synuclein fibrillization and Parkinson’s disease? Nat. Cell Biol. 2000, 2, E115−E119. 10.1038/35017124

8 Tuttle, M. D.; Comellas, G.; Nieuwkoop, A. J.; Covell, D. J.; Brthold, D. A.; Kloepper, K. D.; Courtney, J. M.; Kim, J. K.; Barclay, A. M.; Kendall, A.;, et al. Solid-state NMR structure of a pathogenic fibril of full-length human α-synuclein. Nat. Struct. Mol. Biol. 2016, 23, 409–415. 10.1038/nsmb.3194

9 Li, B.; Ge, P.; Murray, K. A.; Sheth, P.; Zhang, M.; Nair, G.; Sawaya, M. R.; Shin, W. S.; Boyer, D. R.; Ye, S.;, et al. Cryo-EM of full-length α-synuclein reveals fibril polymorphs with a common structural kernel. Nat. Commun. 2018, 9, 3609. 10.1038/s41467-018-05971-2

10 Li, Y.; Zhao, C.; Luo, F.; Liu, Z.; Gui, X.; Luo, Z.; Xiang, Z.; Li, D.; Liu, C.; Li, X. Amyloid fibril structure of α-synuclein determined by cryo-electron microscopy. Cell. Res. 2018, 28, 897–903. 10.1038/s41422-018-0075-x

11 Guerrero-Ferreira, R.; Taylor, N. M.; Mona, D.; Ringler, P.; Lauer, M. E.; Riek, R.; Britschgi, M.; Stahlberg, H. Cryo-EM structure of alpha-synuclein fibrils. eLife 2018, 7, e36402. 10.7554/eLife.36402

12 Guerrero-Ferreira, R.; Taylor, N. M.; Arteni, A.-A.; Kumari, P.; Mona, D.; Ringler, P.; Britschgi, M.; Lauer, M. E.; Makky, A.; Verasdonck, J.;, et al. Two new polymorphic structures of human full-length alpha-synuclein fibrils solved by cryo-electron microscopy. eLife 2019, 8, e48907. 10.7554/eLife.48907

13 Shammas, S. L.; Garcia, G. A.; Kumar, S.; Kjaergaard, M.; Horrocks, M. H.; Shivji, N.; Mandelkow, E.; Knowles, T. P. J.; Mandelkow, E.; Klenerman, D. A mechanistic model of tau amyloid aggregation based on direct observation of oligomers. Nat. Commun. 2015, 6, 7025. 10.1038/ncomms8025

14 Czaplewski, C.; Kalinowski, S.; Liwo, A.; Scheraga, H. A. Application of multiplexed replica exchange molecular dynamics to the UNRES force field: Tests with a and α+β proteins. J. Chem. Theory Comput. 2009, 5, 627–640. 10.1021/ct800397z

15 Patel, J. S.; Ytreberg, F. M. Fast calculation of protein-protein binding free energy using umbrella sampling with a coarse-grained model. J. Chem. Theory Comput. 2018, 14, 991–997. 10.1021/acs.jctc.7b00660

16 Liwo, A.; Czaplewski, C.; Pillardy, J.; Scheraga, H. A. Cumulant-based expressions for the multibody terms for the correlation between local and electrostatic interactions in the united-residue force field. J. Chem. Phys. 2001, 115, 2323–2347. 10.1063/1.1383989

17 Maisuradze, G. G.; Senet, P.; Czaplewski, C.; Liwo, A.; Scheraga, H. A. Investigation of protein folding by coarse-grained molecular dynamics with the UNRES force field. J. Phys. Chem. A 2010, 114, 4471–4485. 10.1021/jp9117776

18 Sieradzan, A. K.; Sans-Duno, J.; Lubecka, E. A.; Czaplewski, C.; Lipska, A. G.; Leszczynski, H.; Ocetkiewicz, K. M.; Proficz, J.; Czarnul, P.; Krawczyk, H.;, et al. Optimization of parallel implementation of UNRES package for coarse-grained simulations to treat large proteins. J. Comput. Chem. 2023, 44, 602–625. 10.1002/jcc.27026

19 Maisuradze, G. G.; Liwo, A.; Scheraga, H. A. Principal component analysis for protein folding dynamics. J. Mol. Biol. 2009, 385, 312–329. 10.1016/j.jmb.2008.10.018

20 Maisuradze, G. G.; Liwo, A.; Ołdziej, S.; Scheraga, H. A. Evidence, from simulations, of a single state with residual native structure at the thermal denaturation midpoint of a small globular protein. J. Am. Chem. Soc. 2010, 132, 9444–9452. 10.1021/ja1031503

21 Maisuradze, G. G.; Liwo, A.; Senet, P.; Scheraga, H.A. Local vs global motions in protein folding. J. Chem. Theory Comput. 2013, 9, 2907–2921. 10.1021/ct4001558

22 Kachlishvili, K.; Maisuradze, G. G.; Martin, O. A.; Liwo, A.; Vila, J. A.; Scheraga, H. A. Accounting for a mirror-image conformation as a subtle effect in protein folding. Proc. Natl. Acad. Sci. U.S.A. 2014, 111, 8458–8463. 10.1073/pnas.1407837111

23 Zhou, R.; Maisuradze, G. G.; Suñol, D.; Todorovski, T.; Macias, M. J.; Xiao, Y.; Scheraga, H. A., Czaplewski, C.; Liwo, A. Folding kinetics of WW domains with the united residue force field for bridging microscopic motions and experimental measurements. Proc. Natl. Acad. Sci. U.S.A. 2014, 111, 18243–18248. 10.1073/pnas.1420914111

24 Maisuradze, G. G.; Medina, J.; Kachlishvili, K.; Krupa, P.; Mozolewska, M.A.; Martin-Malpartida, P.; Maisuradze, L.; Macias, M.; Scheraga, H. A. Preventing fibril formation of a protein by selective mutation. Proc. Natl. Acad. Sci. U.S.A. 2015, 112, 13549–13554. 10.1073/pnas.1518298112

25 Rojas, A.; Maisuradze, N.; Kachlishvili, K.; Scheraga, H. A.; Maisuradze, G. G. Elucidating important sites and the mechanism for amyloid fibril formation by coarse-grained molecular dynamics. ACS Chem. Neurosci. 2017, 8, 201–209. 10.1021/acschemneuro.6b00331

26 Rojas, A.V.; Maisuradze, G. G.; Scheraga, H. A. Dependence, of the formation of tau and Aβ peptide mixed aggregates on the secondary structure of the N-terminal region of Aβ. J. Phys. Chem. B 2018, 122, 7049–7056. 10.1021/acs.jpcb.8b04647

27 Sole-Domenech, S.; Rojas, A. V.; Maisuradze, G. G.; Scheraga, H. A.; Lobel, P.; Maxfield, F. R. Lysosomal enzyme tripeptidyl peptidase 1 destabilizes fibrillar Aβ by multiple endoproteolytic cleavages within the β-sheet domain. Proc. Natl. Acad. Sci. U.S.A. 2018, 115, 1493–1498. 10.1073/pnas.1719808115

28 Xie, H.; Rojas, A.; Maisuradze, G. G.; Khelashvili, G. Mechanistic kinetic model reveals how amyloidogenic hydrophobic patches facilitate the amyloid-β fibril elongation. ACS Chem. Neurosci. 2022, 13, 987–1001. 10.1021/achchemneuro.1c00801

29 Guzzo, A.; Delarue, P.; Rojas, A.; Nicolaï, A.; Maisuradze, G. G.; Senet, P. Missense mutations modify the conformational ensemble of the α-synuclein monomer which exhibits a two-phase characteristic. Front. Mol. Biosci. 2021, 8, 786123. 10.3389/fmolb.2021.786123

30 Guzzo, A.; Delarue, P.; Rojas, A.; Nicolaï, A.; Maisuradze, G. G.; Senet, P. Wild-type α-synuclein and variants occur in different disordered dimers and pre-fibrillar conformations in early stage of aggregation. Front. Mol. Biosci. 2022, 9, 910104. 10.3389/fmolb.2022.910104

31 Yu, H.; Han, W.; Ma, W.; Schulten, K. Transient β-hairpin formation in α-synuclein monomer revealed by coarse-grained molecular dynamics simulation. J. Chem. Phys. 2015, 143, 243142. 10.1063/1.4936910

32 Hoyer, W.; Cherny, D.; Subramaniam, V.; Jovin, T. M. Impact of the acidic C-terminal region comprising amino acids 109-140 on α-synuclein aggregation in vitro. Biochemistry 2004, 43, 16233–16242. 10.1021/bi048453u

33 Trexler, A. J.; Rhoades, E. Single molecule characterization of α-synuclein in aggregation-prone states. Biophys. J. 2010, 99, 3048–3055. 10.1016/j.bpj.2010.08.056

34 Hong, D.-P.; Xiong, W.; Chang, J.-Y.; Jiang, C. The role of the C-terminus of human α-synuclein: Intra-disulfide bonds between the C-terminus and other regions stabilize non-fibrillar monomeric isomers. FEBS Lett. 2011, 585, 561–566. 10.1016/j.febslet.2011.01.009

35 Jain, K.; Ghribi, O.; Delhommelle, J. Folding free-energy landscape of α-synuclein (35-97) via replica exchange molecular dynamics. J. Chem. Inf. Model. 2021, 61, 432–443. 10.1021/acs.jcim.0c01278

36 Khalili, M.; Liwo, A.; Rakowski, F.; Grochowski, P.; Scheraga, H. A. Molecular dynamics with the united-residue model of polypeptide chains. I. Lagrange equations of motion and tests of numerical stability in the microcanonical mode. J. Phys. Chem. B 2005, 109, 13785–13797. 10.1021/jp058008o

37 Khalili, M.; Liwo, A.; Jagielska, A.; Scheraga, H. A. Molecular dynamics with the united-residue model of polypeptide chains. II. Langevin and Berendsen-bath dynamics and tests on model α-helical systems. J. Phys. Chem. B 2005, 109, 13798–13810. 10.1021/jp058007w

38 Fox, N. K.; Brenner, S. E.; Chandonia, J. -M. SCOPe: Structural classification of proteins-extended, integrating SCOP and ASTRAL data and classification of new structures. Nucl. Acids Res. 2014, 42, D304–D309. 10.1093/nar/gkt1240

39 Chandonia, J. -M.; Fox, N. K.; Brenner, S. E. SCOPe: Classification of large macromolecular structures in the structural classification of proteins-extended database. Nucl. Acids Res. 2019, 47, D475–D481. 10.1093/nar/gky1134

40 Kabsch, W.; Sander, C. Dictionary of protein secondary structure: pattern recognition of hydrogen-bonded and geometrical features. Biopolymers 1983, 22, 2577–2637. 10.1002/bip.360221211

41 Grassein, P.; Delarue, P.; Nicolaï, A.; Neiers, F.; Scheraga, H. A.; Maisuradze, G. G.; Senet, P. Curvature and torsion of protein main chain as local order parameters of protein unforlding. J. Phys. Chem. B 2020, 124, 4391–4398. 10.1021/acs.jpcb.0c01230

42 Ramachandran, G. N.; Ramakrishnan, C.; Sasisekharan, V. Stereochemistry of polypeptide chain configurations. J. Mol. Biol. 1963, 7, 95–99. 10.1016/s0022-2836(63)80023-6

43 Doherty, C. P. A.; Ulamec, S. M.; Maya-Martinez, R.; Good, S. C.; Makepeace, J.; Khan, G. N.; van Oosten-Hawle, P.; Radford, S. E.; Brockwell, D. J. A short motif in the N-terminal region of α-synuclein is critical for both aggregation and function. Nat. Struct. Mol. Biol. 2020, 27, 249–259. 10.1038/s41594-020-0384-x

44 Ulamec, S. M.; Maya-Martinez, R.; Byrd, E. J.; Dewison, K. M.; Xu, Y.; Willis, L. F.; Sobott, F.; Heath, G. R.; van Oosten Hawle, P.; Buchman, V. L.;, et al. Single residue modulators of amyloid formation in the N-terminal P1-region of α-synuclein. Nat. Commun. 2022, 13, 4986. 10.1038/s41467-022-32687-1

45 Korneev, A.; Begun, A.; Liubimov, S.; Kachlishvili, K.; Molochkov, A.; Niemi, A. J.; Maisuradze, G. G. Exploring structural flexibility and stability of α-synuclein by the Landau-Ginzburg-Wilson approach. J. Phys. Chem. B 2022, 126, 6878–6890. 10.1021/acs.jpcb.2c04651

46 Krokhotin, A.; Liwo, A.; Maisuradze, G. G.; Niemi, A. J.; Scheraga, H. A. Kinks, loops, and protein folding, with protein A as an example. J. Chem. Phys. 2014, 140, 025101. 10.1063/1.4855735

47 Feig, M.; Karanicolas, J.; Brooks, C. L. 3^rd^. MMTSB Tool Set: enhanced sampling and multiscale modeling methods for applications in structural biology. J. Mol. Graph. Model. 2004, 22, 377–395. 10.1016/j.jmgm.2003.12.005

48 Hollingsworth, S. A.; Karplus, P. A. A fresh look at the Ramachandran plot and the occurrence of standard structures in proteins. BioMol. Concepts 2010, 1, 271–283. 10.1515/bmc.2010.022

49 Khammari, A.; Arab, S. S.; Ejtehadi, M. R. The hot sites of α-synuclein in amyloid fibril formation. Sci. Rep. 2020, 10, 12175. 10.1038/s41598-020-68887-2

50 Vasili, E.; Konig, A.; Al-Azzani, M.; Bosbach, C.; Gatzemeier, L. M.; Thom, S.; Chegao, A.; Mirands, H. V.; Steinem, C.; Erskine, D.; et al. Glycation of alpha-synuclein enhances aggregation and neuroinflammatory responces. npj Parkinsons Dis. 2025, 11, 307. 10.1038/s41531-025-01159-w

51 Best, R. B.; Zhu, X.; Shim, J.; Lopes, P. E. M.; Mittal, J.; Feig, M.; MacKerell, A. D. Jr. Optimization of the additive CHARMM all-atom protein force field targeting improved sampling of the backbone ϕ, ψ and side-chain χ_1_ and χ_2_ dihedral angles. J. Chem. Theory Comput. 2012, 8, 3257–3273. 10.1021/ct300400x

52 Hornak, V.; Abel, R.; Okur, A.; Strockbine, B.; Roitberg, A.; Simmerling, C. Comparison of multiple Amber force fields and development of improved protein backbone parameters. *Proteins: Struct., Funct.*, Genet. 2006, 65, 712−725. 10.1002/prot.21123

53 Guzzo, A. Effets des mutations sur la structure et la dimérisation de l’alpha-synucléine et son implication dans la maladie de Parkinson : réponses apportées par la dynamique moléculaire à gros grains. Ph.D. Dissertation, Université Bourgogne Franche-Comte, Dijon, 2022. 10.70675/0849285az2731z415cza277z0e56728f62bc

54 Nguyen, V. T. T.; Ulamec, S. M.; Brockwell, D. J.; Radford, S. E.; Hall, C. K. In silico peptide self-assebly reveals the importance of N-terminal motifs and the inhibition mechanism of the mutation L38M in α-synuclein fibrillation. Protein Sci. 2026, 35, e70530. 10.1002/pro.70530

55 Rojas, A.; Liwo, A.; Browne, D.; Scheraga, H. A. Mechanism of fiber assembly: treatment of Aβ-peptide aggregation with a coarse-grained united-residue force field. J. Mol. Biol. 2010, 404, 537–552. 10.1016/j.jmb.2010.09.057

