## Supplemental Figures for "Examining the Atypical Structure Adopted by the 37-51 Segment of α-synuclein in the Early Stage of Aggregation"

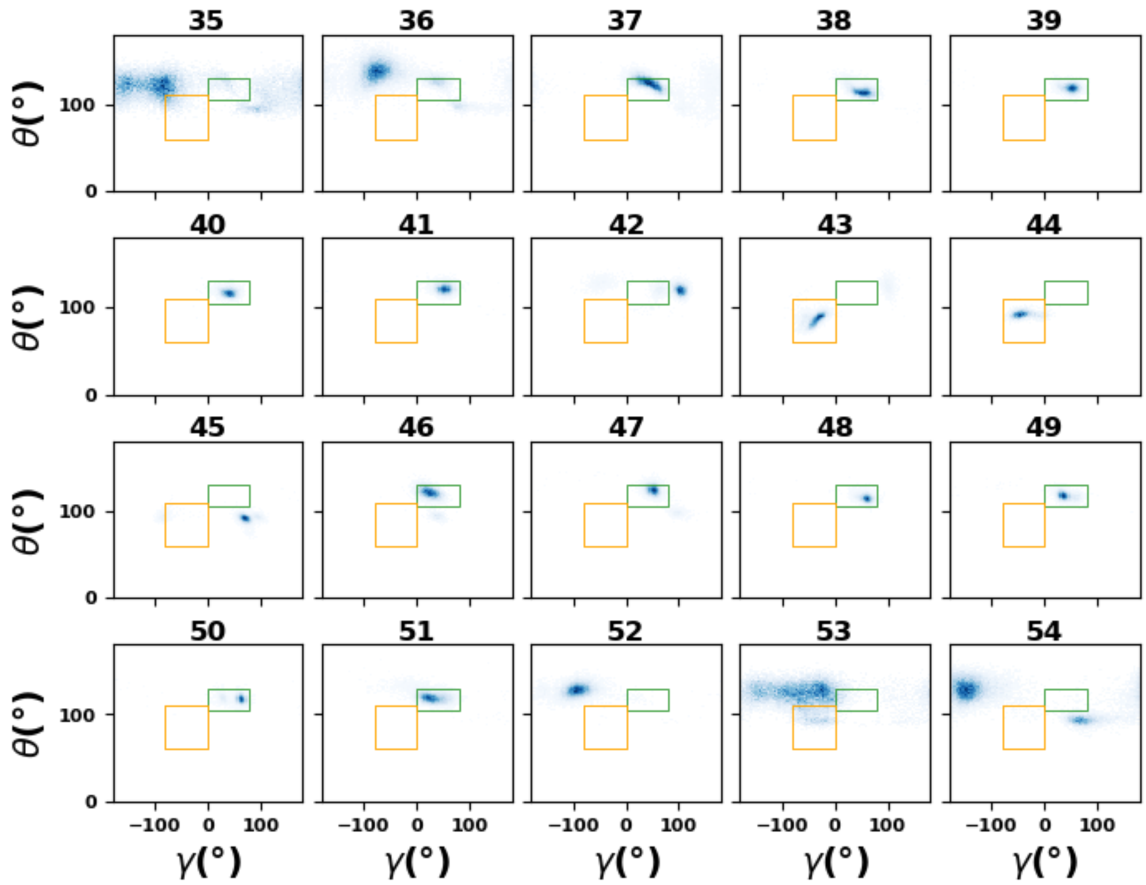


Figure S1. Heatmap of θ/γ angles for each residue in the braid region (residues 35-54). The body and turn regions are highlighted with green and orange rectangles, respectively.

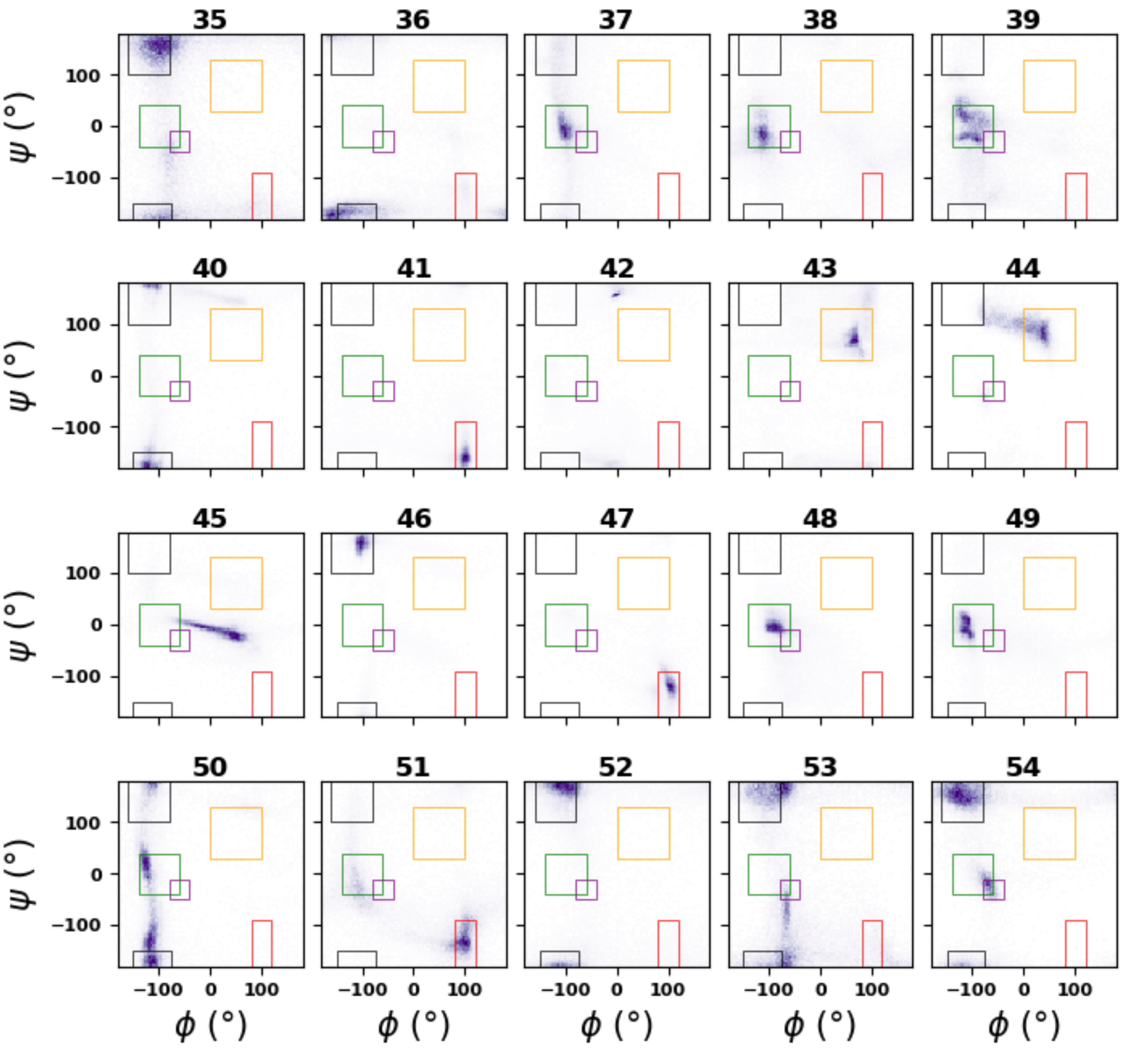


Figure S2. Heatmap of ψ/ϕ angles for each residue in the braid region (residues 35-54). The right-handed α-helix, β-sheet, body, turn, and ε regions are highlithed with purple, black, green, orange, and red rectangles, respectively.
